# *Tempus et Locus*: a tool for extracting precisely dated viral sequences from GenBank, and its application to the phylogenetics of primate erythroparvovirus 1 (B19V)

**DOI:** 10.1101/061697

**Authors:** Alice R. Carter, Derek Gatherer

## Abstract

The presence of data in the “collection_date” field of a GenBank sequence record is of great assistance in the use of that sequence for Bayesian phylogenetics using “tip-dating”. We present *Tempus et Locus* (TeL), a tool for extracting such sequences from a GenBank-formatted sequence database. TeL shows that 60% of viral sequences in GenBank have collection date fields, but that this varies considerably between species. Primate erythroparvovirus 1 (human parvovirus B19 or B19V) has only 40% of its sequences dated, of which only 112 are of more than 4 kb. 100 of these are from B19V sub-genotype 1a and were collected from a mere 6 studies conducted in 5 countries between 2002 and 2013. Nevertheless, Bayesian phylogenetic analysis of this limited set gives a date for the common ancestor of sub-genotype 1a in 1990 (95% HPD 1981-1996) which is in reasonable agreement with estimates of previous studies where collection dates have been assembled by more laborious methods of literature search and direct enquiries to sequence submitters. We conclude that although collection dates should become standard for all future GenBank submissions of virus sequences, accurate dating of ancestors is possible with even a small number of sequences if sampling information is high quality.

## Introduction

Bayesian phylogenetics has become one of the principal technical disciplines of modern virology (Drummond et al., 2006). Packages such as Mr Bayes (Huelsenbeck and Ronquist, 2001) and BEAST (Drummond and Rambaut, 2007) have made such analysis possible on a variety of computer architectures, from Unix clusters to domestic PCs, and the BEAUti interactive GUI (Drummond et al., 2012) packaged with BEAST has made configuration of Bayesian parameter optimizations into virtually a point-and-click process. Bayesian analysis has been applied to deep phylogeny of DNA viruses (Ehlers et al., 2010; Leendertz et al., 2009), using Yule models of speciation and palaentological information to specify priors on the dates of internal nodes of trees, but its principal application has been to the recent evolution of fast-evolving RNA viruses, generating insights into the origins and spread of many viruses, including several of global clinical importance such as HIV-1, hepatitis C and influenza A (Faria et al., 2014; McNaughton et al., 2015; Worobey et al., 2014).

For such viruses, where there is no guide to the dates of internal nodes on the tree, calibration of substitution rate is performed via sampling dates of the sequences used, referred to as “tip-dating” (Rambaut, 2000). Where the geographical locations of samples are also known, phylogenetic analysis of the ancestry of the virus may be complemented by phylogeographic analysis of its spread (Lemey et al., 2010). This combination of probabilistic evolutionary analysis in space and time made a crucial contribution to the tracking and control of the west African outbreak of Zaire ebolavirus in 2013-2015 (Woolhouse et al., 2015). However, although the value of publication of accurate time and place of sampling of viral genomes has now been amply demonstrated by the Zaire ebolavirus epidemic, previously the practice was highly variable in its implementation. In the field of influenza, it was adopted early, although frequently time of collection was only published at the resolution of calendar year. In other viruses, it has often been neglected.

GenBank records include a dedicated “FEATURES” field - “source” - for sampling information. Figure 1 shows the “source” field for an HIV-1 sequence. The precise tip-date is found under the “collection_date” subfield and the location of sampling under “country”. Together, these provide the information needed for the inclusion of this sequence in a phylogenetic/phylogeographic analysis.

**Figure 1:**
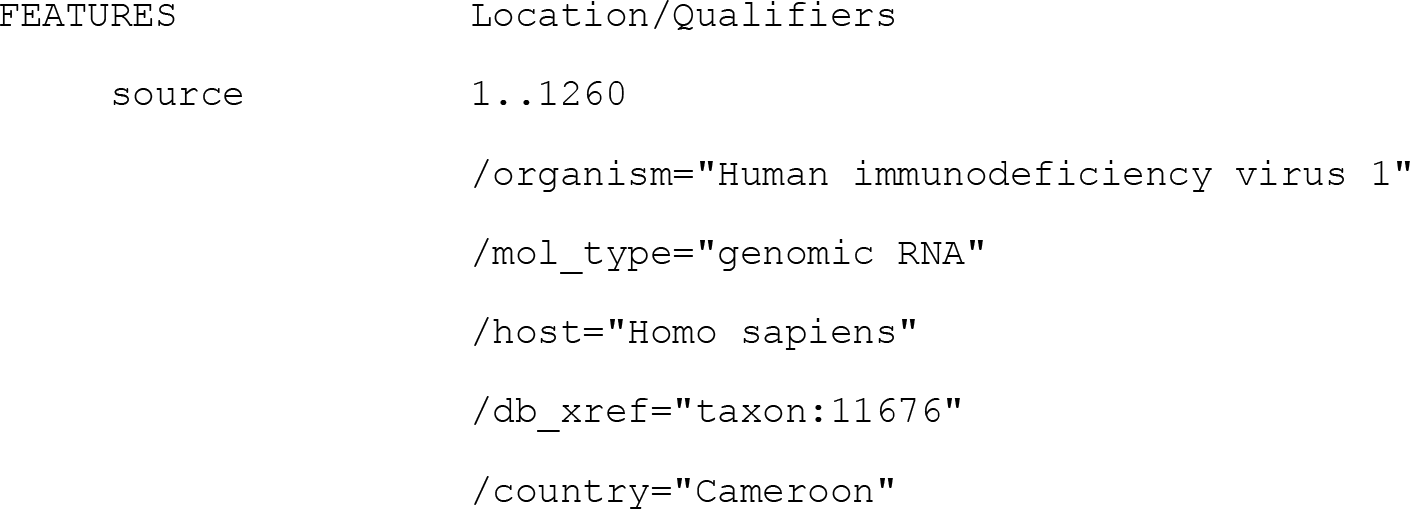
Example source field within FEATURES for GenBank accession KR229865

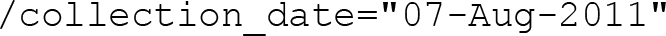

However, assembly of a dataset including this information by direct query on the GenBank website is not straightforward. A GenBank query allows filtering only on “species”, “molecule types”, “source databases”, “genetic compartments”, “sequence length”, “release date” and “revision date”. The second last of these does not correspond to “collection_date”, although it does provide a latest possible tip-date. No field on location of sampling is available for filtering. Consequently, there is no ready means of rapidly extracting tip-dated or tip-located sequences from the GenBank website in large quantities. The exception to this general rule are the viruses included in the Virus Variation Resource (Brister et al., 2014) section of GenBank, which currently includes influenza, dengue, West Nile, MERS and Ebola viruses, and which permits customization of the FASTA defline to include sampling information where available. Until Virus Variation Resource is extended to all virus families, or equivalent data extraction facilities are more generally provided on the main GenBank webpage, this paper describes a tool, *Tempus et Locus* (TeL), based on BioPerl (Stajich, 2007), that aims to fill that gap. Some statistics on the relative abundance of such TeL-positive sequences in the viral section of GenBank are derived and discussed.

Application of TeL is also illustrated with reference to Primate erythroparvovirus 1 (genus Erythroparvovirus; sub-family Parvovirinae; family Parvoviridae), commonly known as human parvovirus B19 (B19V) (Cotmore et al., 2014). B19V is of clinical interest as the cause of the childhood disease erythema infectiosum (Pattison, 1987) and a wide range of more serious complications (Page et al., 2015), and has been the subject of previous phylogenetic analyses (Hubschen et al., 2009; Servant et al., 2002), including some using Bayesian methods (Barros de Freitas et al., 2007; Shackelton and Holmes, 2006). In many of those papers, tip-dates on trees are given, although it is often not completely transparent how they are derived. In some cases, a “collection_date” field appears to have been used, in others it appears that some more complicated deduction has been made - for instance the paper related to the sequence may have time and place of sampling buried in its Materials & Methods section. In others still, one simply surmises that an estimate has been made based on publication date.

Such combinations of guesswork and lengthy reading of the literature are clearly unsatisfactory. A tool such as TeL, described here, would enable the rapid extraction of accurately tip-dated sequences from a GenBank-formatted database. However, given the sparsity of such annotation in some viruses, also discussed below, it begs the question as to whether such a process would thin the dataset to the extent of failure of any subsequent Bayesian phylogenetic analysis. By comparison of Bayesian analysis of a TeL-generated B19V dataset and previous phylogenetic analysis of B19V in the literature, we aim to answer that question.

### Methods

#### Sequences

Nucleic acid sequences from the viral section of GenBank were downloaded on 16^th^ November 2015. Tip-dated sequences were extracted using TeL.

#### Software

TeL is available from doi:10.17635/lancaster/researchdata/52 and requires BioPerl (Stajich, 2007). TeL was developed and tested on Bio-Linux 8.0 (Field et al., 2006) only, but in principle should operate on any system where BioPerl can be installed, including PCs. TeL was used here for extraction of any sequences containing a “collection_date” field, but can also be configured to collect sequences with location of sampling, or both time and location. Sequence alignments of extracted B19V sequences were performed using Muscle (Edgar, 2004) in MEGA (Kumar et al., 2008: http://www.megasoftware.net) and preliminary neighbour joining trees (Saitou and Nei, 1987) constructed. Clock-like behaviour in sequence evolution on those trees was checked using TempEst (Rambaut et al., 2016). Bayesian phylogenetic analysis was performed in BEAST v.1.8.2 (Drummond and Rambaut, 2007: http://tree.bio.ed.ac.uk/software/beast/).

#### Optimization of Bayesian parameters

The strategy for determining the best Bayesian model was as follows. A coalescent constant size tree prior was fixed, and strict clock, relaxed lognormal clock and relaxed exponential clock were run for 10 million iterations in BEAST. The nucleotide substitution model was determined by prior analysis in MEGA to be Tamura 3-parameter (T93+G+I) (Tamura, 1992), and was not subsequently varied. The best of the three clock models was then determined using the Akaike Information Criterion Markov (AICM) tool in Tracer (from the BEAST package), and further runs initiated using that best clock model and expansion growth, logistic growth, exponential growth and a Bayesian skyline. A burn-in of 25% of all trees was used prior to deriving the time to most recent common ancestor (t-MRCA) and its 95% highest probability density (95% HPD) from the consensus tree.

### Results

#### TeL-positive sequences in GenBank viral section

A total of 1,869,455 viral nucleic acid sequences were obtained from GenBank. TeL extracted 1,125,163 with collection dates, just over 60%. Total processing time was 114 mins on an Intel(R) Xeon(R) CPU E5-2690 v2 @ 3.00GHz running Bio-Linux 8.0. The largest single species in terms of dated entries is influenza A with 329,951, closely followed by HIV-1 with 305,608. Hepatitis C is a distant third, on 56,630. Table 1 shows the distribution of the top 14 TeL-positive species across GenBank. Some manual curation of the statistics was required to produce Table 1 as species annotation is not always consistent, for instance dengue virus serotypes are listed separately, as are influenza A subtypes etc.

**Table 1:**
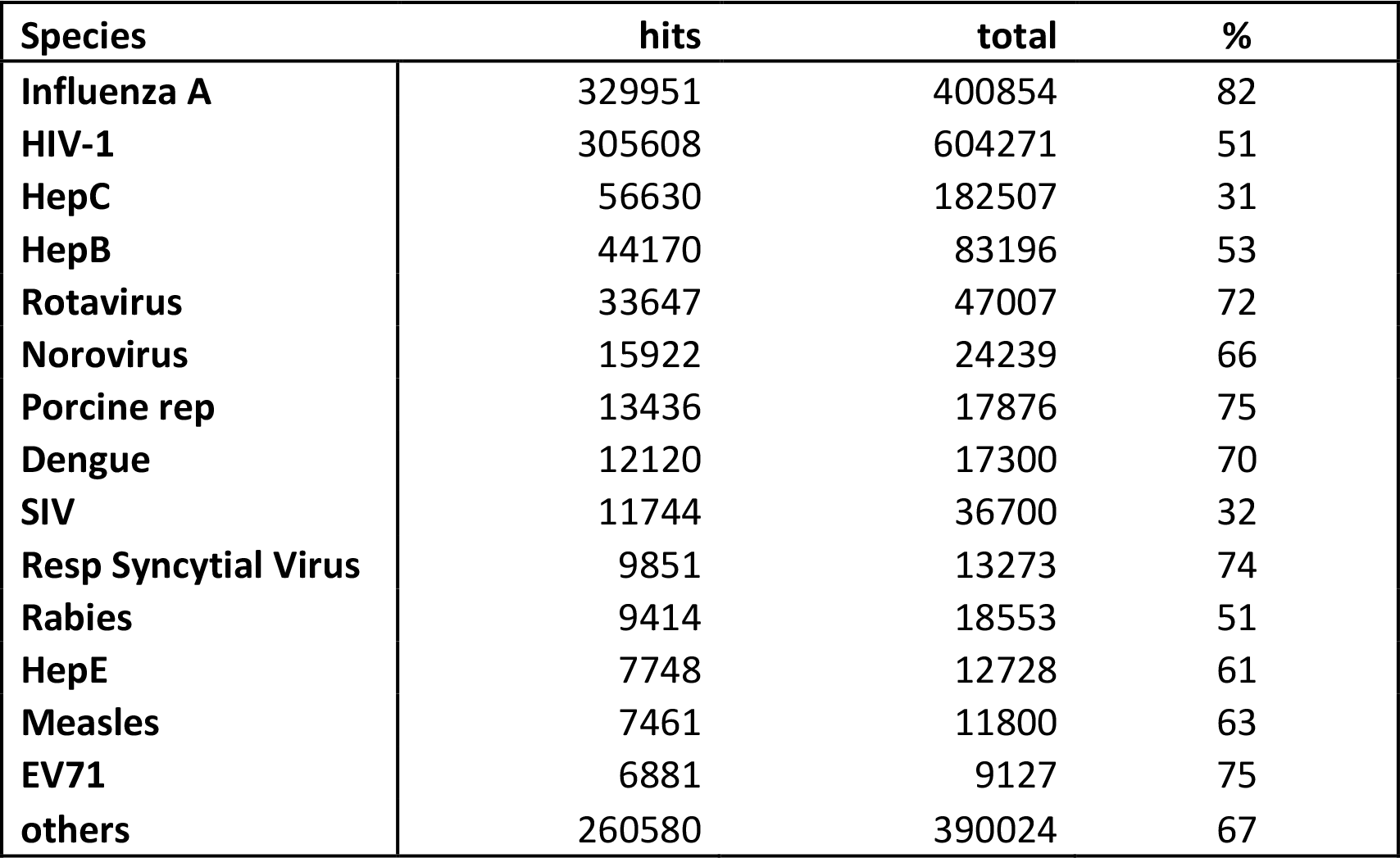
The top 14 species in terms of their total TeL hits for collection date. **Hits**: total TeL-positive sequences per species; **total**: total GenBank sequences per species; %: percentage TeL-positive per species.

Against the overall average of 60%, there is a relative enrichment of TeL-positive sequences in the following species within the top 14 as assessed by number of TeL hits: influenza A, rotavirus, norovirus, porcine reproductive and respiratory syndrome virus, dengue virus, respiratory syncytial virus, hepatitis E, measles and EV71. Hepatitis C and SIV sequences are by contrast poorly dated, with only around half the GenBank average for TeL-positivity.

#### TeL-positive sequences from B19V

A total of 2723 B19V nucleic acid sequences were found in GenBank, of which 1077 are TeL-positive-at just under 40% this is considerably lower than the GenBank average of 60%. 112 of these are over 4kb in length and were selected for further Bayesian phylogenetic analysis as representative of full-length or near-full-length genomes. The earliest of these was collected in Belgium on 22^nd^ February 2002 and the most recent consists of a group of seven sequences collected in the USA in 2013. This gives a window of a mere 11 years in which to calibrate the molecular clock. The first B19V sequence of over 4kb to be released was in 1986 (Shade et al., 1986), and the earliest genome sequenced in retrospect is from 1973 (Blundell et al., 1987). The obvious question therefore arises as to whether or not insistence on accurate tip-date as provided by “collection_date” field is too stringent a criterion.

Furthermore, the distribution of TeL-positive sequences was predominantly B19V genotype 1 (Servant et al., 2002), at 101 out of 112. Sequences from genotypes 2 and 3 - four and seven genomes respectively-were therefore not analysed further. One further sequence (KC013303-da Costa et al., 2013) was annotated in GenBank as a recombinant and also removed, leaving 100 sequences. Table 2 classifies these remaining Tel-positive B19V genotype 1 genomes, which are derived from a mere six studies conducted in three European countries, the USA and Brazil, and all of which are subtype 1a. Only two of these datasets appear to have any associated literature (da Costa et al., 2013; Molenaar-de Backer et al., 2012).

**Table 2:** Sources of the Tel-positive B19V genotype 1a genomes. ‡The range KC013305- KC013351 given for the Brazilian sequences includes only KC0133+suffix: 05, 08, 12-14, 16, 21, 22, 24, 25, 27, 29, 31-33, 38, 40, 43, 44, 46, 51.

| Accession range | Geographical origin | Collection date range | Reference |
| --- | --- | --- | --- |
| FJ591158 | USA | 2008 | Chen,Z. et al (unpublished) |
| FN598217 | Belgium | 22 Feb 2002 | Op de Beeck,A. et al (unpublished) |
| JN211121-JN211185 | Netherlands | March 2003-August 2009 | (Molenaar-de Backer et al., 2012) |
| KC013305-KC013351<sup>‡</sup> | Brazil | 2007-2010 | (da Costa et al., 2013) |
| KM393163-KM393169 | USA | 2013 | Qiu,Y. et al (unpublished) |
| KR005640-KR005644 | Serbia | 2009-2012 | Stamenkovic,G.G. et al (unpublished) |

#### Clock-like evolution of B19V genotype 1a

Figure 2 presents the TempEst analysis of the neighbour joining tree of the B19V genotype TeL-positive genotype 1a sequences, plotting root-to-tip distance on a neighbour-joining tree against sampling date. MEGA determined the best substitution model to be T92+G+I and the value of the alpha parameter in the gamma distribution to be 0.794. TempEst’s best fitting line (R=0.50) crosses the x-axis at 1982 and determines the best-fitting tree under the assumption of a strict clock to be a mid-point rooted one (Figure 3), suggesting that sub-genotypes 1a1 and 1a2 both evolved from a common ancestor, rather than either emerging as a sub-clade from the other. No genotype 1b sequences are TeL-positive and so, like genotypes 2 and 3, remain unanalysed here.

**Figure 2:**
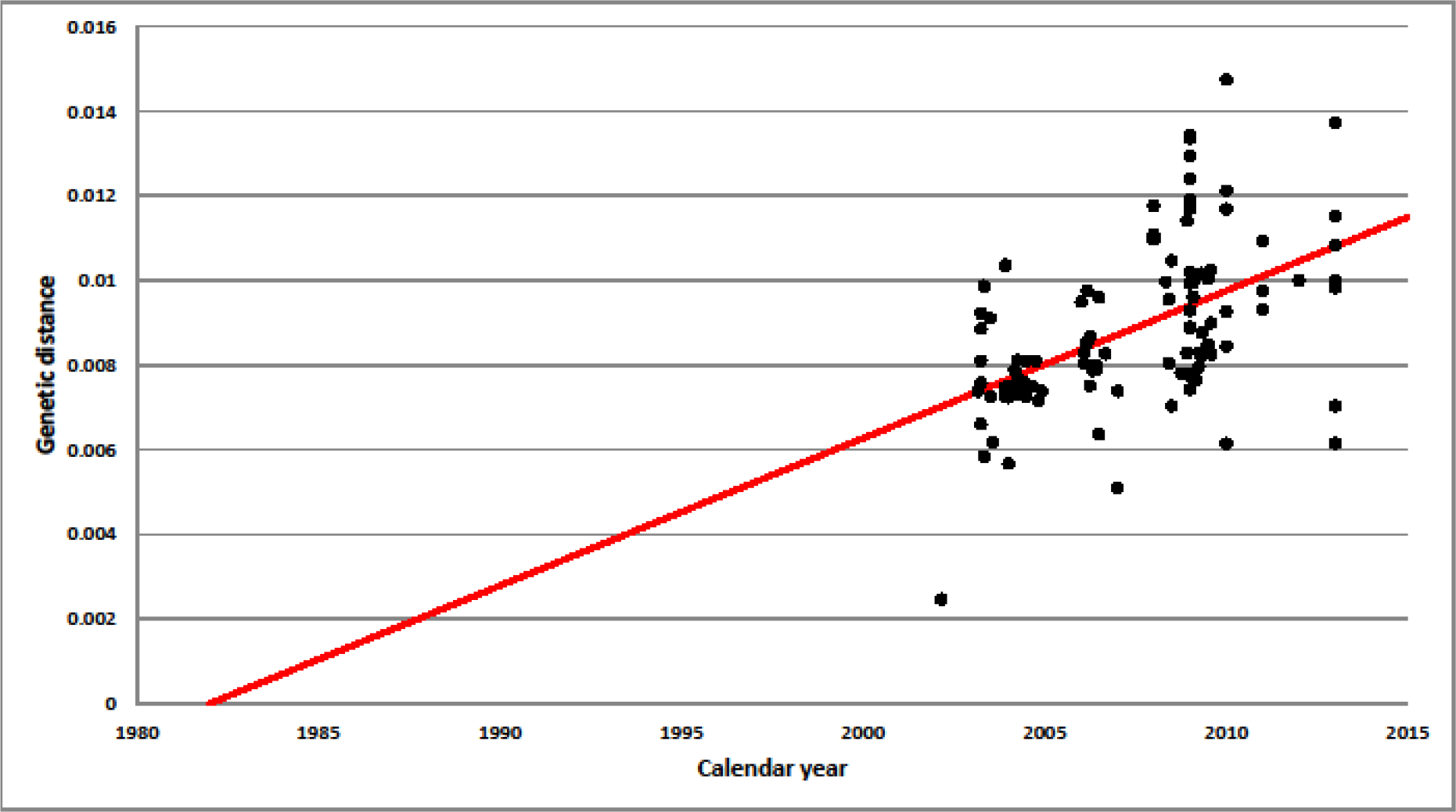
TempEst plot of genetic distance in substitutions per site from the mid-point root of a neighbour-joining tree against sampling date. Red: Best-fitting line, correlation coefficient 0.50.

**Figure 3:**
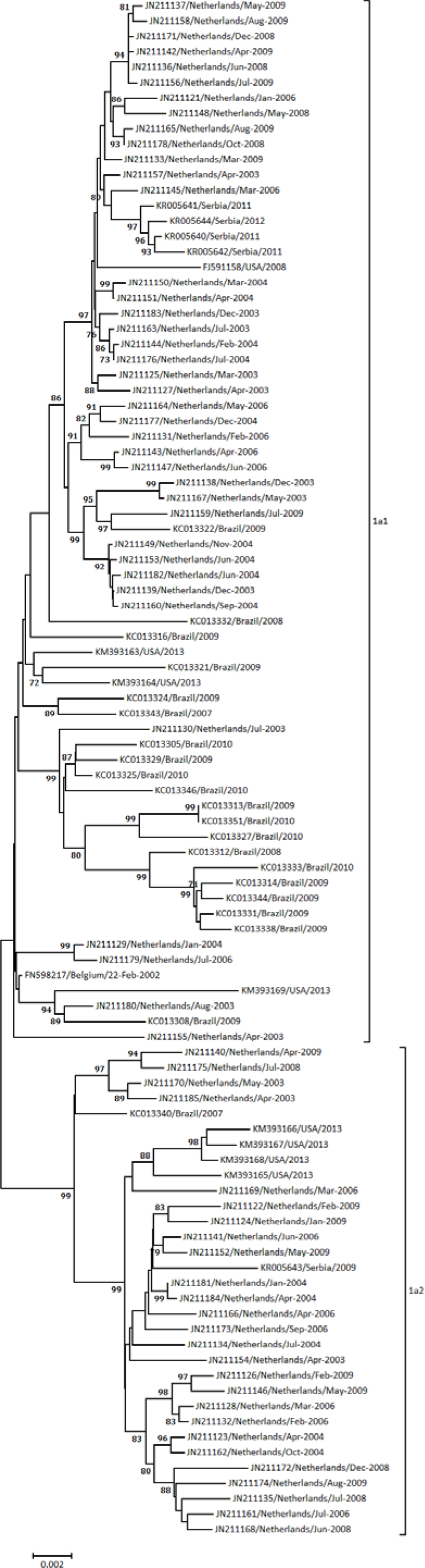
Mid-point-rooted neighbour joining tree corresponding to Figure 2, consistent with sub-genotypes 1a1 and 1a2 being clades descended from a putative common 1a ancestor. Bootstrap values over 70 are shown.

#### Bayesian phylogenetic analysis of B19V genotype 1a

The relaxed exponential clock was first determined to be the best clock model under the assumption of a coalescent constant size tree prior. Using the relaxed exponential clock, exponential growth was then determined to be the best tree prior. Table 3 shows the z-scores for model comparison. The time of the most recent common ancestor (t-MRCA) was placed at 23 years before the most recent dated sequence (2013) or 1990, with 95% HPD of 17 to 32 years or 1981-1996, using the best model (relaxed exponential clock with exponential growth). This is somewhat later than the x-axis intersection of the TempEst analysis at 1982 (Figure 1), although that year does lie just inside the 95% HPD. BEAST analysis over all the models generally produces a mean t-MRCA in the range 21-27 years before 2013, except for the relaxed exponential clock with a constant population size when the t-MRCA is at 42 years and has a much wider 95% HPD. Because of this discrepancy, that parameter set was run five times on two different servers, both with and without the BEAGLE library, but the results were consistent across all runs. We have no explanation for this discrepancy, but disregard it since model comparison shows it to be an inferior scenario.

**Table 3:**
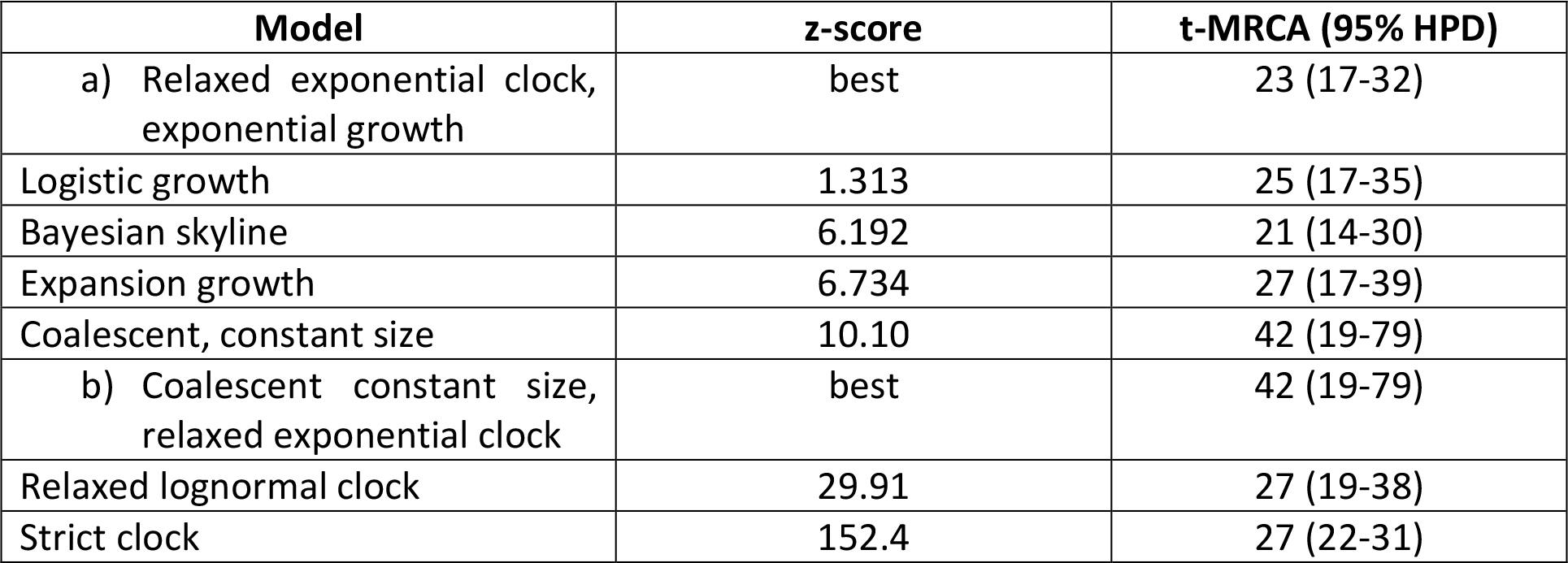
Akaike Information Criterion Markov (AICM) (Baele et al., 2012) z-scores for candidate models of growth and clock. A) Comparison of exponential growth model with other tree priors, all run with relaxed exponential clock; B) Comparison of relaxed exponential clock with other clock models, all run under coalescent constant size population model. The relaxed exponential clock is a markedly better fit to the data than other clock models (B) and exponential growth is better than other growth models, although not significantly so for logistic growth (A).

| Model | z-score | t-MRCA (95% HPD) |
| --- | --- | --- |
| a) Relaxed exponential clock, exponential growth | best | 23 (17-32) |
| Logistic growth | 1.313 | 25 (17-35) |
| Bayesian skyline | 6.192 | 21 (14-30) |
| Expansion growth | 6.734 | 27 (17-39) |
| Coalescent, constant size | 10.10 | 42 (19-79) |
| b) Coalescent constant size, relaxed exponential clock | best | 42 (19-79) |
| Relaxed lognormal clock | 29.91 | 27 (19-38) |
| Strict clock | 152.4 | 27 (22-31) |

### Discussion

#### The relatively sparsity of collection-dated genomes for B19V and other viruses

TeL only selected those B19V sequences with “collection_date” fields, of which only those of greater than 4kb length were selected for Bayesian analysis. For B19V, this means that no genotype 1b sequences, and only a mere handful of genotype 2 and 3, are available, and the Bayesian analysis is confined to genotype 1a. B19V has “collection_date” information for only 40% of its GenBank sequences, well below the overall average of 60%. One might be tempted to speculate that this might relate to B19V’s relatively low clinical profile but Table 1 shows that HIV-1, Hepatitis B, Hepatitis C and rabies are also all below the 60% average. The inclusion of “collection_date” fields is therefore to be encouraged in the future. The figure of 40% Tel-positive sequences would be even smaller for B19V if it were not for one recent large-scale study in the Netherlands (Molenaar-de Backer et al., 2012).

#### Estimation of the date of the common ancestor of B19V genotype 1a

Barros de Freitas et al. (2007) previously performed a Bayesian phylogenetic analysis of 139 B19V sequences, using a 477 bp segment, and Figure 1 of their paper shows the common ancestor of genotype 1a at approximately 14 years after their common ancestor of all B19V sequences. They assumed, following Shackelton and Holmes (2006), a root on the oldest sample, from 1973 (Blundell et al., 1987). The scale on their Figure 1 is not entirely clear: the legend seems to imply an origin at the time of the oldest sequence which would place the ancestor of genotype 1a at 14 years after 1973, or 1987. This is within three years of our estimate using our best model (1990 - Table 3A). However, it is also possible to interpret their scale as having year 35 as corresponding to their most recent sequence, or 2005. This would then place the root at 1970 and ancestor of genotype 1a at 1984, towards the earlier tail of our 95% HPD and closer to the date of 1982 suggested by our TempEst best-fitting line on root-to-tip genetic distance (Figure 2). Barros de Freitas and colleagues (2007) also derived logistic growth as the best demographic model, which we find to be non-significantly inferior to our best model (Table 3A). Their clock model is specified as relaxed, but they do not state if it is relaxed exponential or relaxed logarithmic.

An earlier Bayesian analysis using 24 sequences of over 4kb - thus more directly comparable to the current study - did not find any significant differences between relaxed and strict clocks (Shackelton and Holmes, 2006: Table 1 of that paper) in terms of the derived mean substitution rates. This is compatible with our finding (Table 3) that all models suggest a mean t-MRCA for the common ancestor for genotype 1a around 21-27 years before 2013-with the exception of the anomalous result for the relaxed exponential clock and coalescent constant size, for which we have no explanation. The dataset of Shackelton and Holmes (2006) ranged from 1973 to 2001 and thus has no overlap with the sequences studied here. Unfortunately their study places no dates on any of the internal nodes of their tree, so we have no point of comparison for the ancestor of genotype 1a.

Other phylogenetic studies of B19V have ignored the question of internal tree dates. For instance the paper of Hubschen et al. (2009), uses a set of 166 sequences for which nearly 1kb of sequence data was derived. Unfortunately, none of these appear to have collection dates. To take two examples for illustrative purposes, GenBank accession FN295632, representing the Bishkek06-155 B19V strain, has no “collection_date” field, and can only be tip-dated by Table 1 of Hubschen et al. (2009) in which “2006-2007” is given. Likewise FN295577, representing the BGR-Deniss strain, has a “2005-2007” date given in the same Table.

#### Dates of older common ancestors

The date of the common ancestor between genotypes also remains to be conclusively demonstrated. The two previous Bayesian studies place the root of the entire B19V tree on the 1973 sequence. The practice of rooting trees for fast evolving RNA viruses on the oldest sequence has had demonstrable success recently in clarifying the phylogenetic position of the 2013-2015 Makona strain of Zaire ebolavirus (Dudas and Rambaut, 2014), doing so principally by arranging the topology of the tree in such a way as to maximise clock-like behaviour, which may then be judged using TempEst, as presented here in Figure 2 (and Figure 5 of Dudas and Rambaut, 2014). Shackelton and Holmes (2006) present a similar plot in Figure 1 of their paper which might suggest a radiation of B19V into its current 3 genotypes in the last half century. However, it is not clear how well this hypothesis sits with observations that genotype 2 is only found in individuals born before 1973, or that genotype 3 - the African genotype-has greater diversity than the other two, which Servant-Delmas et al. (2010) suggest is indicative of a deeper genetic history in Africa. There is therefore scope for a deeper study of the origins of B19V. This, however, will require a larger number of accurately tip-dated sequences from genotypes 2 and 3.

#### Conclusions

It is hoped that *Tempus et Locus* (TeL) will prove a useful tool for the quick extraction of accurately tip-dated and tip-located sequences from GenBank. We have shown that such information is highly variable between viral families. We also show that even when it is relatively sparse and unequally distributed over time, such as in B19V, Bayesian trees can be produced in BEAST which are reasonably consonant with earlier studies. We note in passing that the molecular phylogenetics of B19V has a few issues that remain to be clarified, in particular the date of the most recent common ancestor of all current genotypes.

## Data Access Statement

*Tempus et Locus* code, manual, input data, derived data, alignments used for neighbour-joining and Bayesian analysis, and all MEGA and BEAST outputs are all available from: doi:http://10.17635/lancaster/researchdata/52

## Acknowledgement

We thank reviewers at *Virus Evolution* for their comments on the original submitted manuscript.

